# Ghrelin is related to lower brain reward activation during touch

**DOI:** 10.1101/2022.05.10.491384

**Authors:** D.M. Pfabigan, E.R. Frogner, E. Schéle, P. M. Thorsby, B. S Skålhegg, S. L. Dickson, U. Sailer

**Author notes:** Corresponding author: Daniela M. Pfabigan, Jonas Lies vei 91, 5009 Bergen, mail.

## Abstract

The gut hormone ghrelin drives food motivation and increases food intake, but it is also involved in the anticipation of and response to rewards other than food. This pre-registered study investigated how naturally varying ghrelin concentrations affect the processing of touch as a social reward in humans.

Sixty-seven volunteers received slow caressing touch (so-called CT-targeted touch) as a social reward and control touch on their shins during 3T functional imaging on two test days. On one occasion participants were fasted and on another they received a meal. On each occasion plasma ghrelin was measured at three time points.

All touch was rated as more pleasant after the meal, but there was no association between ghrelin concentrations and pleasantness. CT-targeted touch was rated as most pleasant and activated somatosensory and reward networks (whole-brain). A region-of-interest in the right medial orbitofrontal cortex (mOFC) showed lower activation during all touch the higher ghrelin concentrations were. During CT-targeted touch, a larger satiety response (ghrelin decrease after the meal) was associated with higher mOFC activation, and this OFC activation was associated with higher experienced pleasantness.

Overall, higher ghrelin concentrations appear to be related to lower reward value of touch. Ghrelin may reduce the value of social stimuli, such as touch, to promote food search and intake in a state of low energy. This suggests that the role of ghrelin goes beyond assigning value to food reward.

## 1. Introduction

Ghrelin is a hormone that is secreted by cells in the stomach and regulates appetite (Kojima et al. 1999). As it particularly increases the motivation to eat foods high in sugar and fat (Perello 2010; Egecioglu et al. 2010), even in the absence of hunger (Skibicka et al. 2012), it seems to act specifically on reward aspects of eating. Besides food, ghrelin’s effect also extends to other types of rewards. For example, its administration makes mice consume more alcohol (Jerlhag et al. 2009) and raises the motivation to self-administer heroin in rats (Maric et al. 2012). Ghrelin also increased the preference for social interaction in one group of rats, whereas a ghrelin antagonist decreased it (Schéle et al., 2020). On the other hand, the suppression of ghrelin signalling reduced alcohol consumption in rats and mice (e.g.; Jerlhag et al., 2009; Zallar et al., 2019), and reduced the rewarding effects of amphetamine and nicotine in mice (Jerlhag et al. 2010; Jerlhag and Engel 2011) and their sexual motivation (Egecioglu et al., 2016). In humans, ghrelin concentrations are associated with the amount of craving in individuals with alcohol dependence (Leggio et al. 2012), with the rated pleasantness of odours (Trellakis et al. 2011; but see also (Pfabigan et al., 2022)) and reward system activation in response to food pictures (Malik et al. 2008). Following ghrelin administration, participants showed less brain activity during the anticipation of monetary losses than following administration of saline, and they also valued receiving a monetary reward sooner rather than later (Pietrzak et al., 2023).

This widespread effect of ghrelin may occur because different types of rewards operate within the same reward system (Berridge and Kringelbach 2015). The rewarding effects arise via the activity of the mesolimbic dopamine system which originates in the ventral tegmental area (VTA). That ghrelin engages the mesolimbic reward system is suggested by studies in rodents in which ghrelin delivery (either peripherally or to the VTA) increases dopamine release in the nucleus accumbens (Jerlhag et al., 2009). Consistent with this, ghrelin delivery to this site increases food motivation, reflected by increased lever-pressing for sucrose in an operant responding paradigm (Skibicka et al., 2012, 2011). In this way, ghrelin promotes food intake by increasing the incentive and rewarding responses to food cues (e.g.; Abizaid et al., 2006; Kawahara et al., 2013). Because ghrelin regulates dopaminergic function in the VTA (Abizaid 2009; Skibicka and Dickson 2011), it may have a more general role of modulating neural function in the reward system. As such, ghrelin may not only modify the response to and motivation for food and alcohol in humans, but also to non-food rewards, for instance social rewards derived from interacting with other individuals. Therefore, the rationale of the current study was to investigate if ghrelin may modulate the response to social reward in the form of touch.

Slow stroking is considered a social reward, because it occurs in social interactions, is typically experienced as pleasant (Essick et al., 2010; e.g., Kass-Iliyya et al., 2017; Morrison et al., 2011; Triscoli et al., 2013), and participants work to obtain it (Løseth et al., 2019; Perini et al., 2015). Stroking velocities between 3 and 10 cm/s best stimulate the so-called C-tactile (CT) nerve afferents (Ackerley et al., 2014; Löken et al., 2009), which are assumed to transmit emotionally relevant touch in social settings. In line with this assumption, CT-targeted touch activates brain areas involved in social processing such as the superior temporal gyrus (Sailer et al., 2016) and sulcus (Björnsdotter et al., 2014; Perini et al., 2021), in addition to the posterior insula and secondary somatosensory cortex (McCabe et al., 2008). Activation in reward areas such as orbitofrontal cortex (Grabenhorst et al., 2010; Sailer et al., 2016) and nucleus accumbens (Kreuder et al., 2017; McCabe et al., 2008) has also been reported occasionally.

To investigate if ghrelin is related to touch reward, the neural activation during touch and its subjective experience were assessed in two different sessions in 67 healthy volunteers, in a cross-over within-subject fast-and-feed approach. In one of the sessions, participants arrived fasted and continued fasting throughout to increase ghrelin concentrations. In the other session, the same participants received a standardised meal with the aim to decrease ghrelin concentrations. This procedure allowed us to assess the effects of naturally varying intra-individual ghrelin concentrations. Our pre-registered bi-directional hypothesis was that being in a fasted state with high ghrelin concentrations is either associated with increased touch reward similar to the previously reported reward-enhancing effects of ghrelin (Egecioglu et al., 2016; Jerlhag et al., 2009; Jerlhag and Engel, 2011; Maric et al., 2012; Schéle et al., 2020), or is associated with decreased touch reward, because ghrelin can also condition a place aversion (Schéle et al., 2017). A further pre-registered hypothesis was that the difference in ROI brain activation for CT-targeted touch in the sessions with and without the meal would be associated with differences in ghrelin concentrations in those sessions.

## 2. Methods

The study was preregistered on the Open Science Forum in October 2019 (https://osf.io/f9rkq), deviations from the pre-registration are clearly stated in the manuscript. Data collection took place in 2019 and 2020.

### 2.1 Participants

The initial sample consisted of 68 participants, 47 men (mean age: 27.8, SD: 7.8), 20 women (mean age: 31.9, SD: 10), and one participant who did not wish to disclose their gender (age: 30). One male participant dropped out after the first blood sample because of circulation problems. Seven participants were not available for the second session that was delayed due to an intervening Covid-19 lockdown. Sixty participants, 42 men, 17 women, and one undisclosed individual completed both sessions. Potential participants (between 18 and 55 years old) were recruited via flyers and advertisements in social media. Further inclusion criteria were sufficient Norwegian language skills, a body mass index (BMI) between 18 and 29.9 kg/m^2^, normal or corrected to normal vision, and fulfilment of MRI safety criteria. In addition, participants had no history of eating disorders, diabetes, gastrointestinal surgery, not taking any drugs affecting gastrointestinal function. It was a prerequisite that female participants did not take hormonal contraceptives. All female participants’ measurements were scheduled in the first week of their menstrual cycle (days 1-8) to avoid additional variance in reward processing (Bedwell et al., 2019; Dreher et al., 2007; Pletzer et al., 2019), and to reduce potential effects of gonadal hormones on ghrelin concentrations (Börchers et al., 2022). Further demographic information (handedness, educational attainment, self-reported ancestry) can be found in Supplementary Materials (section 2.1).

A priori power analysis (Westfall, 2016) was calculated for pleasantness ratings and a medium effect for the interaction between touch velocity and nutritional state (*d*=0.45). A sample size of 66 participants was recommended to reach 80% power, but the Covid-19 pandemic did not allow us to reach the planned sample size – see Supplementary Materials (section 2.1). Deviating from the power analysis in the pre-registration, we applied single trial analyses for the behavioural data with linear mixed models instead of a means-based ANOVA design. Thus, the current behavioural analyses reached a power of 80% for a small effect of the touch velocity x nutritional state interaction (d=0.3, replicants=5 (i.e, 5 repetitions per touch velocity)) with 36 participants.

Participants were reimbursed for their participation with universal gift cards (∼ EUR 150 per person for both sessions, of which ∼EUR 50 for the first one). During a typical four to five-hour session, participants conducted further experimental tasks which are (Pfabigan et al., 2022; Sailer et al., 2023) and will be presented elsewhere (i.e., tasks on pain experience and sweet taste perception).Participants gave written informed consent prior to the experiment, which was approved by the Regional Ethics Committee (REK South-East B, project 26699).

### 2.2 Procedure

The same participants were invited twice to the laboratory. On two separate test days, all participants arrived at the laboratory after having fasted for 6 hours. In order to compare differing endogenous ghrelin concentrations, on one of these visits participants received a standard liquid meal which is known to decrease ghrelin concentrations (“liquid-meal” session). On the other visit they received no meal, i.e. participants remained fasted with high ghrelin concentrations throughout the whole session (“no-meal” session). Session order was pseudorandomized between participants; the two test sessions were by median four days apart (range 1-85). Overall, 65 liquid-meal and 62 no-meal sessions were conducted. Participants were instructed to drink no more than 1 litre of water and fast in the six hours before experiment start, which was usually at 3 pm. To motivate participants’ compliance to fasting, they were told that blood glucose concentrations will be assessed via a pin prick test at the beginning of each test session (using an Accu-Check® Aviva device (Roche Diagnostics Norge AS) and test stripes, which assess gut hormones in blood plasma and cortisol in saliva (T0/Baseline). A bioelectrical impedance analysis measurement (seca mBCA 515, seca gmbh & co. kg., Germany) was conducted during the second test session (after the first blood sample) to assess body composition variables such as BMI, fat mass and adipose visceral tissue, which are out of scope of the current report and were not considered in any statistical model as covariates. In the liquid-meal session, participants consumed 300 ml of raspberry flavoured, fat-free, fermented milk (Biola®, Tine BA) and 300 ml of chocolate milk (Sjokomelk, Tine BA) within 15 minutes after the first fasting blood sample. In the no-meal session, participants were offered those two beverages at the end of the experiment.

Participants provided ratings on subjective bodily states and their affective state after the liquid meal and a similar time point in the no meal session, thus most likely reflecting their experience at timepoint T1 (see Supplementary Materials, section 2.2. on a detailed description of the rating questions). Afterwards, participants were instructed in the subsequent experimental tasks, and trained them on a laptop. In the no-meal session, participants were additionally presented with an odour probe after task instructions to assess their ability to identify a food odour and to increase hunger feelings and ghrelin concentrations (Peris-Sampedro et al., 2021). We used peanut butter for this purposes in all participants but one (for whom a banana odour Sniffin’ stick (Hummel et al., 1997) was used due to a nut allergy). Next, the second blood and saliva samples were collected (T1), approximately one hour after the first ones. Then, participants walked over to the imaging facility, located at Oslo University Hospital (10-min walking distance). In the liquid-meal session, participants ate a banana shortly before scanning to keep their stomachs filled and ghrelin concentrations lowered.

Before scanning, participants were made familiar with the response devices at the scanner and completed two training trials with them. The time between the second blood sample (T1) and the touch task was approximately 50 minutes. In the liquid-meal session, the average time between the provided meal and the touch task was approximately 86 minutes. In the no-meal session, the minimum time between the participants’ last meal and the touch task was 8 hours and 10 minutes. After scanning, participants returned to the Institute of Basic Medical Sciences, where the third blood and saliva samples were collected (T2). Figure 1 illustrates the timing of the two experimental sessions.

**Figure 1.**
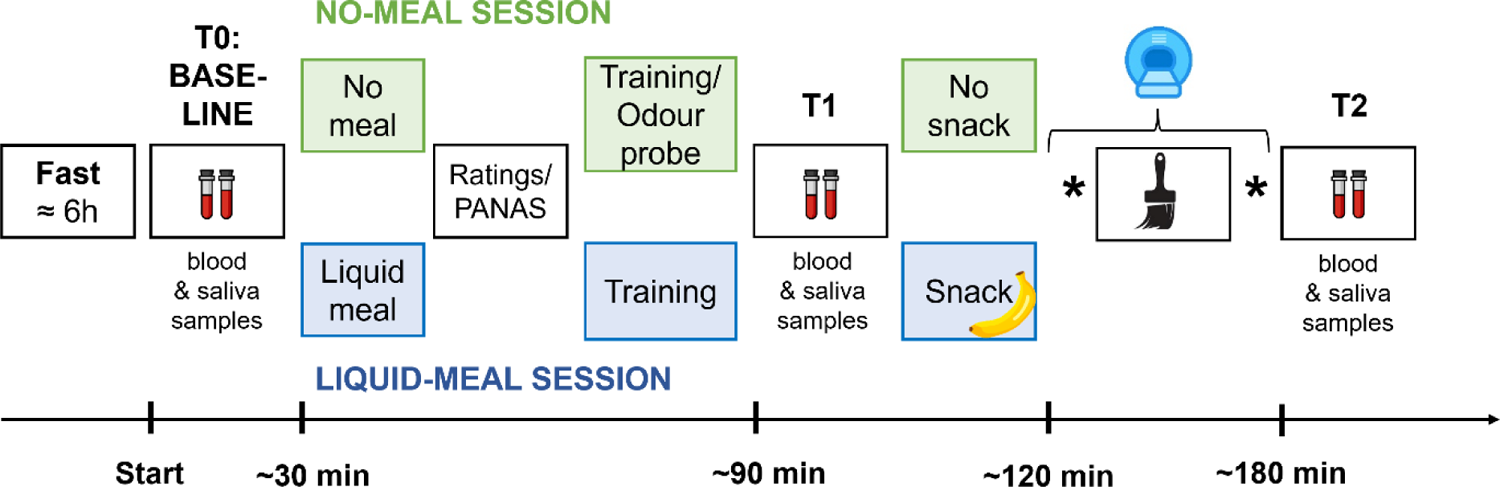
Sequence of experimental procedures. All participants arrived fasted at the laboratory, where a baseline assessment of blood, saliva, and blood glucose was performed first (time point T0/Baseline). Specifics for the no-meal session are illustrated in green colour, specifics for the liquid-meal session in blue colour. After the liquid-meal/no-meal, participants rated their current bodily and affective states. Subsequently, participants were instructed and trained on the upcoming experimental tasks. They were presented with an odour probe in the no-meal session only. Collection of blood and saliva samples after liquid-meal/no-meal followed (T1). Shortly before scanning, a snack was provided in the liquid-meal session only. Stars represent further experimental tasks conducted in the scanner before and after the touch task (brush symbol). After the scanning session, the last blood and saliva samples of the experiment were collected (T2).

### 2.3 Ghrelin analysis

Blood was collected in 2 ml EDTA tubes that were prepared with 100µl protease inhibitor (Pefabloc® SC Plus, Merck KGaA, Germany). These tubes were centrifuged immediately after venepuncture for 15 min at 4⁰C and 3200g. Plasma samples were then aliquoted and stabilized with HCl before storage at −80⁰C in in-house freezer facilities. Ghrelin samples were analysed at the Hormone Laboratory at Oslo University Hospital (Oslo, Norway). Active (acylated) ghrelin concentrations were determined using the EZGRA-88K kit (Merck, Germany) in duplicates (total analytical CV at 488 pg/ml 12%).

### 2.4 Touch task

Participants were placed in the scanner with their right shin exposed while the experimenter stood next to them with a brush. The touch task consisted of one run with 15 trials, with five trials for each of three touch velocities: very slow (0.3 cm/sec), CT-targeted (3 cm/sec), and fast (30 cm/sec). Presentation order of the velocities was randomized across participants. The experimenter brushed the participants’ right shin for 15 seconds per trial, using a 75 mm wide goat-hair brush, always from the knee towards the foot. Stimuli were applied over a length of 10 cm with an approximate force of 0.4 N. All experimenters were trained in applying the correct velocity and force of stroking. Brushing was guided by a visual warning cue via a monitor that was visible from inside the scanning room, and a sliding bar that moved at a set pace while stroking. After each trial, participants rated subjective pleasantness and intensity of touch by answering the questions “*How did you experience the touch?*” and “*How intense did you experience the touch?*” on two visual analogue scales (VAS) with the anchors unpleasant/not intense (coded as 0 for subsequent analyses) and pleasant/intense (coded as 100). Participants had 12 sec to give a response; otherwise, they were presented with a reminder to respond faster on the next trial. The subsequent inter-stimulus interval depicted a white fixation cross on black background. Its duration was jittered between 25 and 27 sec with a uniform distribution (500 ms intervals). The duration of the two VAS scales was subtracted from these values to guarantee a fixed timing for the overall task, which lasted approx. 11 minutes. Stimulus presentation was controlled by E-Prime 2.0/3.0 (Psychology Software Tools, Inc., Sharpsburg, PA). Figure 2 depicts the trial sequence and timing of the touch task.

**Figure 2:**
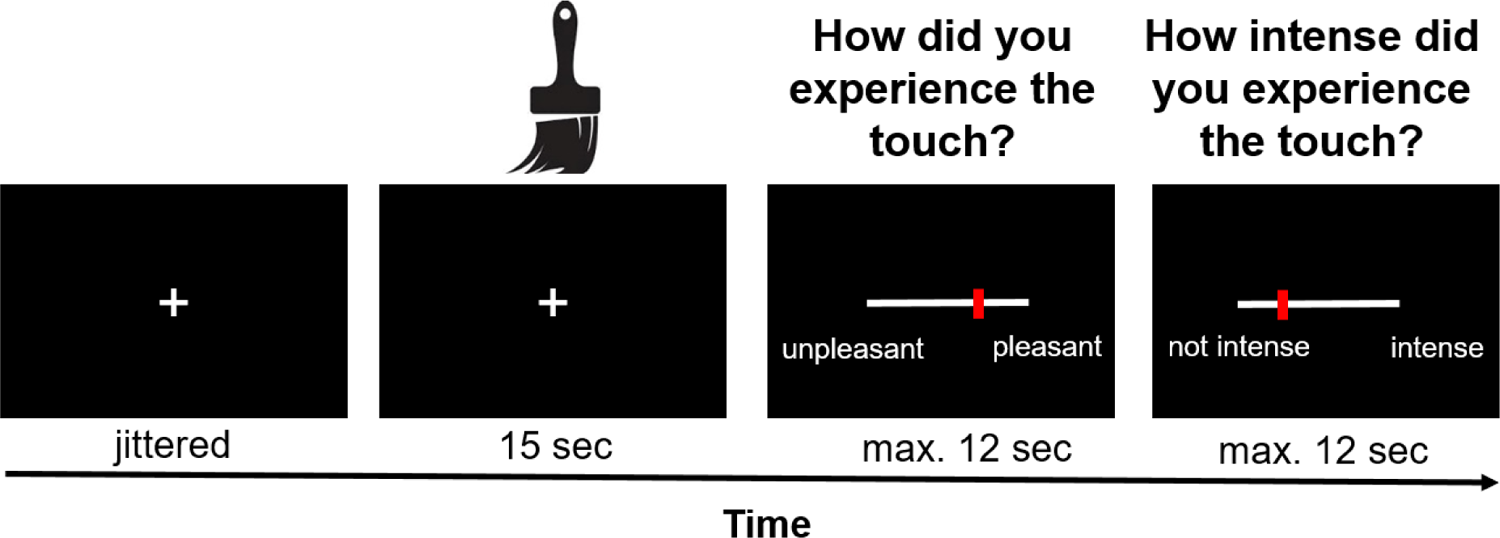
Timing of one touch task trial. After a jittered inter-stimulus interval, the experimenter applied the respective touch stimulation on participants’ right shins for 15 sec. Subsequently, participants rated pleasantness and intensity of the previous touch on two VAS scales via *ResponseGrips* (Nordic Neurolab, Norway).

### 2.5 fMRI data acquisition and analysis

Functional and anatomical MR images were acquired with a 3T Philips Ingenia MRI scanner (Philips Healthcare, Best, NL). For functional imaging, a 32-channel SENSE head-coil was used applying an echo planar imaging (EPI) sequence with the following parameters: voxel size = 3 mm, repetition time (TR) = 2500ms, echo time (TE) = 30ms, flip angle = 90⁰, FOV = 240×240×120, 40 slices, interleaved without gap. After scanning of five dummy scans, 267 functional images were acquired. For the high-resolution anatomical image, a T1-weighted 3D MP-RAGE sequence was applied with the following parameters: TR/TE = 5.2/2.3 ms, flip angle = 8⁰, voxel size = 1 x 1 x 1 mm, FOV = 184×256×256; scan duration: approx. 5 min.

Data pre-processing was carried out with SPM12 (FIL Group, UC London, UK) separately for each scanning session and task. After conversion from dicom to nifti format with MRIcroGL 1.2.20190902++ (Rorden and Brett, 2000), individual brain anatomy files were anonymized with the SPM12 function *De-face images* by deleting image information of facial features. This was not mentioned in the pre-registration, but required by the local ethics board. Default algorithms and parameters were used for slice time correction (to the first slice), motion correction (referenced to the mean image) and unwarping, spatial normalization to MNI (Montreal Neurological Institute) stereotactic space, and spatial smoothing (6 mm FWHM Gaussian kernel). We applied a 4 mm threshold for excessive head movements, which none of the analysed participants exceeded. Moreover, the ArtRepair toolbox (https://cibsr.stanford.edu/tools/human-brain-project/artrepair-software.html) was used to identify artefact-afflicted scanning sessions. Due to an unexpected interaction between the current EPI sequence and a movement-correction setting used during everyday clinical routine at Oslo University Hospital, the data from 15 participants were not usable for whole brain analyses because they showed non-correctable signal distortions in either the liquid-meal or the no-meal session.

Following pre-processing, data were analysed using (1) a whole brain, (2) a region-of-interest (ROI), and (3) a correlation approach. Single-subject analysis (first-level) was performed based on the GLM framework in SPM12 by modelling the three touch velocities and the VAS scales as regressors. Additionally, six realignment parameters were added as nuisance regressors to the model. After estimation, three contrasts were computed for each participant to be included in the group analysis: the three touch velocity conditions were modelled against the implicit baseline (fixation cross). Further details on the first-level analysis can be found in section 2.5, Supplementary Materials.

Group analysis of the whole brain analysis (second-level) implemented a full-factorial model with touch velocity (very slow, CT-targeted, fast) and nutritional state (liquid-meal, no-meal) as repeated factors; 45 artefact-free datasets were available for this analysis. Results are presented at a cluster-level family-wise error (FWE) corrected threshold of *p*<0.05 (starting threshold *p*<0.001 uncorrected).

For the ROI analysis, 60 liquid-meal and 51 no-meal test sessions were available for analysis. Pre-registered ROIs included left posterior insula, left and right secondary somatosensory cortex (SII), right superior temporal gyrus (STG) and left and right medial orbitofrontal cortex (mOFC). Exploratory ROIs included left and right anterior insula and left and right ventral striatum. See Supplementary Materials, section 2.5. for literature references, anatomical and functional definition, and hypotheses for the respective ROIs, as well as the implemented Bonferroni correction for ROI analyses. Parameter estimates were extracted with the REX toolbox (http://web.mit.edu/swg/software.htm). Using linear mixed models, we applied a two-stage approach to disentangle (i) effects of nutritional state, and (ii) associations of ghrelin concentrations with brain activation during touch. These analyses were not pre-registered. Mean ROI activation was first modelled as a function of touch velocity (very slow/CT-targeted/fast), nutritional state (liquid-meal/no-meal), and their interaction as fixed effects (including a random intercept for participant, a random slope for nutritional state, and a random slope for touch velocity if model convergence allowed it). Secondly, mean ROI activation was modelled as a function of touch velocity (very slow/CT-targeted/fast), measurement session (1,2), and mean-centred ghrelin concentrations (sample time point T1), and the interaction of touch velocity x ghrelin as fixed effects (including a random intercept for participant and random slopes for measurement session and touch velocity, if model convergence allowed the latter). For all linear mixed models, we used the Satterthwaite method for approximation of degrees of freedom and applied a restricted maximum likelihood estimation for fixed effects. As effect size measures, we report semi-partial R^2^ (Edwards et al., 2008), for which values of 0.02, 0.13, and 0.26 denote small, medium and large effects (Cohen, 1992). Post-hoc tests were Bonferroni-corrected. Model descriptions can be found in section 2.5, Supplementary Materials.

Lastly, as described in the pre-registration, potential associations between differences in CT-targeted touch ROI brain activation for liquid meal vs. no meal and differences in metabolic state measures, in particular ghrelin, were investigated. We calculated Spearman correlations between difference values (liquid-meal minus no-meal) of ghrelin concentrations at T1 (labelled as Δghrelin) and respective difference values of ROI brain activation in right mOFC (as this was the only brain region significantly associated with ghrelin variation; ΔmOFC for liquid-meal minus no-meal). Thus, Δghrelin reflects differences in absolute ghrelin concentrations at T1 in the liquid-meal vs. the no-meal session. We also explored whether respective differences in pleasantness ratings for CT-targeted touch (Δpleasantness for liquid-meal minus no-meal) were associated with differences in right ΔmOFC activation. Statistical analyses of ROI data were performed with jamovi 1.6.23 (“The Jamovi project”).

### 2.6 Behavioural and hormone data analyses

Statistical analyses of behavioural and hormone data were performed with jamovi 1.6.23 (“The Jamovi project”). Blood glucose concentrations at the beginning of the test sessions, affective state ratings (PANAS), and subjective bodily state ratings were compared between the liquid-meal vs. the no-meal sessions with paired t-tests (blood glucose, PANAS positive affect) or Wilcoxon signed ranks tests for non-normally distributed data (PANAS negative affect, ratings on subjective bodily states).

Deviating from the pre-registration, we conducted linear mixed models to analyse hormone data instead of ANOVAs because linear mixed models can deal with missing values and model individual data more accurately (Barr et al., 2013; Gueorguieva and Krystal, 2004). This is important because due to complications when collecting blood samples, ghrelin concentrations were not available in all participants and all measurement time points. Ghrelin and cortisol concentrations were analysed each with a linear mixed model with nutritional state (liquid-meal/no-meal) and the three sample time points (T0/baseline, T1, T2) as fixed factors. The random effects structure included a random intercept for participant and a random slope for nutritional state. Post-hoc tests were Bonferroni-corrected. Cortisol results are reported in Supplementary Materials (section 3.2).

As with the ROI analysis, linear mixed models and a two-stage approach were applied to analyse subjective ratings to disentangle (i) effects of nutritional state, and (ii) associations of ghrelin concentrations with subjective ratings. First, single-trial pleasantness and intensity ratings were modelled as a function of touch velocity (very slow/CT-targeted/fast), nutritional state (liquid-meal/no-meal), the mean-centred trial number, and the interaction of touch velocity x nutritional state as fixed effects (including a random intercept for participant and random slopes for touch velocity and nutritional state). Second, single-trial pleasantness and intensity ratings were modelled as a function of touch velocity (very slow/CT-targeted/fast), measurement session (1,2), mean-centred trial number, and mean-centred ghrelin concentrations (sample time point T1), and the interaction of touch velocity x ghrelin as fixed effects (including a random intercept for participant and random slopes for touch condition and measurement session). Deviating from the pre-registration, we refrained from modelling ghrelin concentrations and nutritional state (liquid-meal/no-meal) as fixed effects in one model because they share variance with each other. Mean-centred trial number was added a factor to these models to account for changes in ratings over time. More details on the statistical models can be found in section 2.4, Supplementary Materials.

Moreover, exploratory control analyses were carried out to assess potential effects of subjectively experienced negative affect at T1 on behavioural and neural responses of the touch task (see Supplementary Materials, sections 2.6 and 3.5). Further, control analyses with participants’ biological sex and age as covariates were conducted (Supplementary Materials, section 3.6).

## 3. Results

Further details of the results can be found in Supplementary Materials (section 3).

### 3.1 Behavioural and hormonal results

#### 3.1.1 Effectiveness of the nutritional state manipulation

Blood glucose concentrations at the beginning of each measurement sessions did not differ significantly (t(59)=-1.86, p=.067, d=-0.24). Subjective feelings of hunger were lower after the liquid-meal than at a similar time point in the no-meal session (W(59)=1049, p<.001, rank biseral correlation=.784). PANAS positive affect ratings did not differ following liquid-meal and no-meal (t(59)=-0.89, p=.379), but PANAS negative affect ratings were slightly higher following no-meal than liquid-meal (W(59)=635, p=.007, rank biseral correlation=.475). Descriptively, this result appeared to be mostly driven by three items (asking for *nervousness*, *irritability*, and *jittery feelings*). See Supplementary Materials (section 3.1, Table S1) for ratings of further bodily states additional to hunger.

Ghrelin concentrations were significantly affected by nutritional state (b=82.1, SE= 21.0, t(59.0)=3.92, p<.001, semi-partial R^2^=0.21), sample time point (T0/baseline vs. T1: b=-203.2, SE=21.0, t(227.1)=−9.67, p<.001, semi-partial R^2^=0.32; T1 vs. T2: b=69.3, SE=18.0, t(226.9)=3.84, p<.001, semi-partial R^2^=0.32), and their interaction (p’s<.001, semi-partial R^2^=0.29). Fasting ghrelin concentrations were comparable at the beginning of both test sessions (T0/Baseline: difference=-50.71, SE=31.1, t(203)=−1.63, p>.999). Similarly, ghrelin concentrations were comparable at the end of both test sessions (T2: difference =-19.6, SE=31.8, t(209)=-0.62, p>.999). At T1, about 30 min after the liquid meal, ghrelin concentrations were significantly decreased by more than 300 pg/ml compared to the same time point when participants did not receive food (difference =316.74, SE=33.0, t(217)=9.59, *p*<.001). See Table 1 and Figure S1 in Supplementary Materials for ghrelin variation, and section 3.2. for cortisol variation.

**Table 1.** Ghrelin concentrations and touch ratings

|  | Liquid-meal |  |  | No-meal |  |  |
| --- | --- | --- | --- | --- | --- | --- |
|  | *M* | *SD* | *n* | *M* | *SD* | *n* |
| **Acylated ghrelin (pg/ml)** |  |  |  |  |  |  |
| *Baseline (T0)* | 906 | 518 | 62 | 851 | 483 | 60 |
| *T1* | 502 | 274 | 57 | 836 | 445 | 52 |
| *T2* | 843 | 467 | 61 | 779 | 403 | 56 |
| **Pleasantness ratings** |  |  |  |  |  |  |
| *Very slow touch* | 57.1 | 12.1 | 65 | 54.3 | 12.1 | 61 |
| *CT-targeted touch* | 68.5 | 14.5 | 65 | 66.5 | 14.7 | 61 |
| *Fast touch* | 63.0 | 15.2 | 65 | 61.9 | 14.0 | 61 |
| **Intensity ratings** |  |  |  |  |  |  |
| *Very slow touch* | 28.8 | 17.8 | 65 | 30.10 | 18.60 | 62 |
| *CT-targeted touch* | 40.4 | 17.1 | 65 | 39.00 | 17.20 | 62 |
| *Fast touch* | 53.7 | 15.8 | 65 | 53.90 | 18.00 | 62 |

#### 3.1.2 Change of pleasantness and intensity ratings of different touch velocities with nutritional state and ghrelin concentrations

To test if nutritional state had an effect on subjective touch experience, pleasantness and intensity ratings were first modelled as a function of touch velocity, nutritional state, and trial number. Pleasantness ratings were higher for CT-targeted than very slow touch (b=-11.86, SE=1.31, t(67.1)=-9.08, p<.001, semi-partial R^2^=0.55), and for CT-targeted than fast touch (b=5.00, SE=1.51, t(67.1)=3.31, p=.002, semi-partial R^2^=0.55). Pleasantness ratings decreased with trial number (b=-0.30, SE=0.06, t(1662.4)=−5.38, p<.001, semi-partial R^2^=0.02), and were higher following the liquid meal than no meal (b=2.27, SE=1.06, t(61.7)=2.14, p=.036, semi-partial R^2^=0.07). Intensity ratings were higher for fast compared to CT-targeted (b=-14.07, SE=2.07, t(66.2)=-6.81, p<.001, semi-partial R^2^=0.59), and for CT-targeted than very slow touch (b=-10.33, SE=1.39, t(65.4)=-7.43, p<.001, semi-partial R^2^=0.59). Intensity ratings increased with trial number (b=0.41, SE=0.06, t(1672.1)=6.83, p<.001, semi-partial R^2^=0.03). Interaction effects between touch velocity and nutritional state did not reach significance (CT-targeted vs. slow touch x nutritional state: p=.078; CT-targeted vs. fast touch x nutritional state p=.052).

Second, in order to find out if ghrelin concentrations were associated with subjective touch experience, pleasantness and intensity ratings were modelled as a function of touch velocity, trial number, measurement session, and ghrelin concentrations. Main effects for touch velocity were similar to the first model. CT-targeted touch was rated as more pleasant than very slow touch (b=-11.76, SE=1.34, t(61.6)=-8.79, p<.001, semi-partial R^2^=0.56), and more pleasant than fast touch (b=4.46, SE=1.60, t(59.0)=2.78, p=.007, semi-partial R^2^=0.56). Pleasantness decreased with trial number (b=-0.30, SE=0.06, t(1425.3)=-4.95, p<.001, semi-partial R^2^=0.02). Ghrelin variation was not associated with pleasantness (p=.225), and neither was measurement session (p=.298). The intensity of CT-targeted touch was rated to be higher than of slow touch (b=-10.34, SE=1.52, t(60.6)=-6.78, p<.001, semi-partial R^2^=0.58), and the intensity of fast touch was rated higher than of CT-targeted touch (b=-14.60, SE=2.23, t(60.7)=-6.56, p<.001, semi-partial R^2^=0.58). Intensity ratings increased with trial number (b=0.34, SE=0.06, t(1430.9)=5.43, p<.001, semi-partial R^2^=0.02). No association with ghrelin concentrations was observed (p=.882), while intensity was in general rated to be higher during the second measurement session than the first one (b=4.03, SE=1.25, t(51.0)=3.23, p=.002, semi-partial R^2^=0.17). Mean rating data are reported in Table 1.

### 3.2 fMRI results

#### 3.2.1 Change of whole-brain brain activation to touch at different velocities with nutritional state (pre-registered analysis)

Brain activation for CT-targeted touch was higher than for very slow and fast touch in a network of brain regions including left somatosensory cortex (SII), left middle to superior temporal gyrus (STG), bilateral putamen extending into left caudate, right precentral and inferior frontal gyrus extending into the insular cortex, bilateral supplementary motor area (SMA) and precentral gyri, and left lateral occipital cortex/middle temporal gyrus. A main effect of nutritional state indicated higher brain activation following the liquid meal in the right inferior frontal gyrus extending into the frontal operculum. Brain activation in the different nutritional states did not significantly interact with any of the touch velocities (see Table 2 and Figure 3).

**Figure 3:**
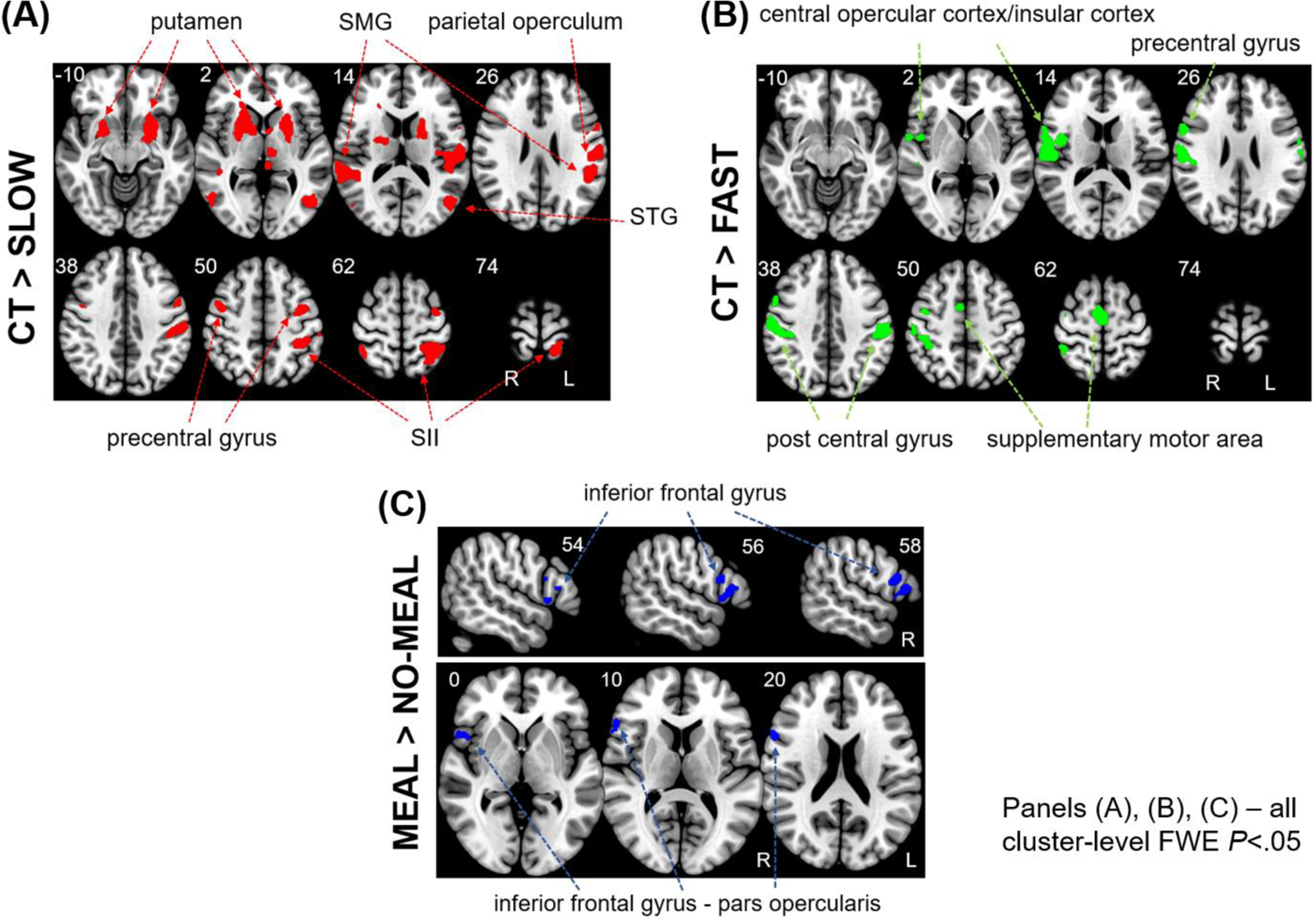
Whole brain results for touch velocity and nutritional state. Panel (A) depicts significant activation clusters for the contrast CT-targeted > very slow touch (in red) in 8 axial slices (Z coordinates reported). Panel (B) depicts significant activation clusters for the contrast CT-targeted > fast touch (in green) in 8 axial slices (Z coordinates reported). Panel (C) depicts the significant activation cluster for the contrast liquid-meal > no-meal (in blue) in 3 sagittal slices in the upper part (right hemisphere, X coordinates reported) and in 3 axial slices in the lower part (Z coordinates reported). All results are presented at a cluster-level FWE corrected level of p<.05.

**Table 2.**
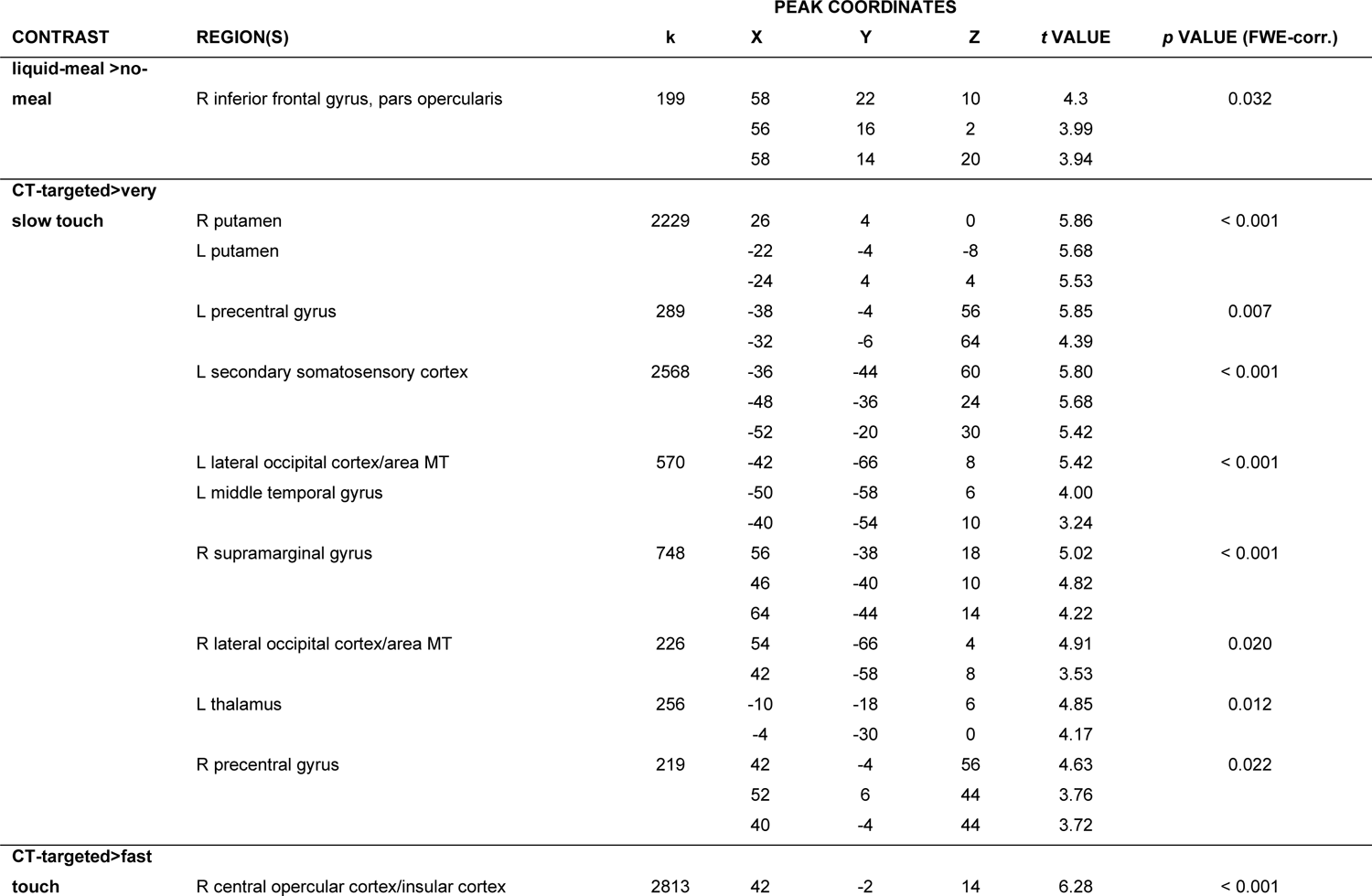
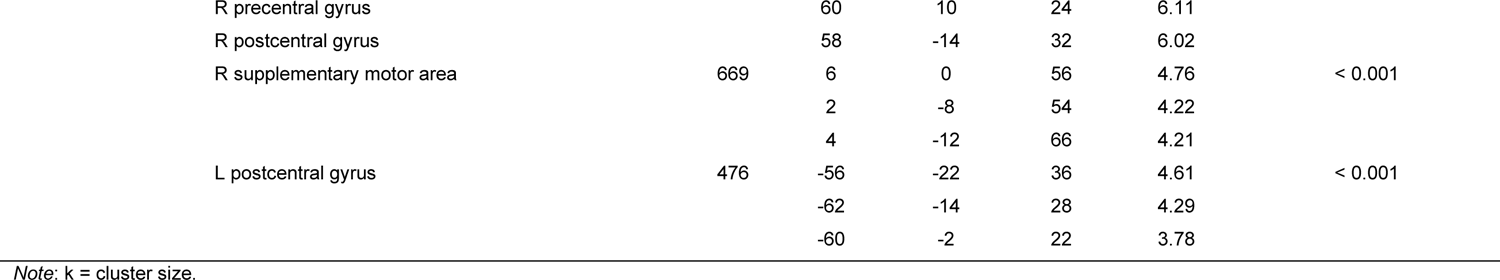
Whole brain results for the main effects of nutritional state and touch velocity

#### 3.2.2 Change of brain activation in different regions-of-interest (ROIs) with touch velocity, nutritional state and ghrelin concentrations (exploratory analyses)

To test if nutritional state changes the brain activation in pre-defined ROIs, we first modelled ROI activation as a function of touch velocity and nutritional state. This showed enhanced activation for CT-targeted than for very slow touch in right STG, bilateral SII and right ventral striatum (VS) (p’s<.004). Activation for CT-targeted than for fast touch was higher in right SII (p<.001). None of the ROIs showed significant differences between liquid meal and no meal (p’s>.055).

To test if ghrelin concentrations were associated with brain activation in those pre-defined ROIs, we modelled ROI activation as a function of touch velocity, measurement session and ghrelin concentrations. Activation was higher for CT-targeted than for very slow touch in right STG, bilateral SII and right VS (p‘s<.007). Activation for CT-targeted was higher than for fast touch in right SII (p<.001). Higher ghrelin concentrations were associated with decreased brain activation in the right medial orbitofrontal cortex (mOFC) irrespective of touch velocity (b=-1.34e^-4^, SE=5.45e^-5^, t(90.2)=−2.46, p=.016, semi-partial R^2^=0.06). Detailed results of these exploratory ROI analyses are reported in Table S3 in Supplementary Materials.

#### 3.2.3 Relationship between nutritional state differences in brain activation during social-affective (CT-targeted) touch and ghrelin concentrations (pre-registered analysis)

Since CT-targeted touch is particularly tuned to social interactions (Ackerley et al., 2014; Löken et al., 2009), we were especially interested in ghrelin’s association with this touch velocity. A significant negative association between Δghrelin at T1 and right ΔmOFC activation was observed (r_s_=-0.412, p=.013, n=36), suggesting that the greater the suppression of ghrelin by the liquid meal compared to no meal (i.e., the satiety response induced by the meal), the higher the brain activation following the liquid meal during CT-targeted touch (compared to no meal; Figure 4B). Furthermore, a significant positive association between right ΔmOFC activation and Δpleasantness was found (r_s_=0.377, p=.012, n=44, Figure 4A), while Δghrelin and Δpleasantness were not associated (p=.914). This means that the higher the brain activation during CT-targeted touch following the liquid meal (compared to no meal), the higher was touch pleasantness.

**Figure 4:**
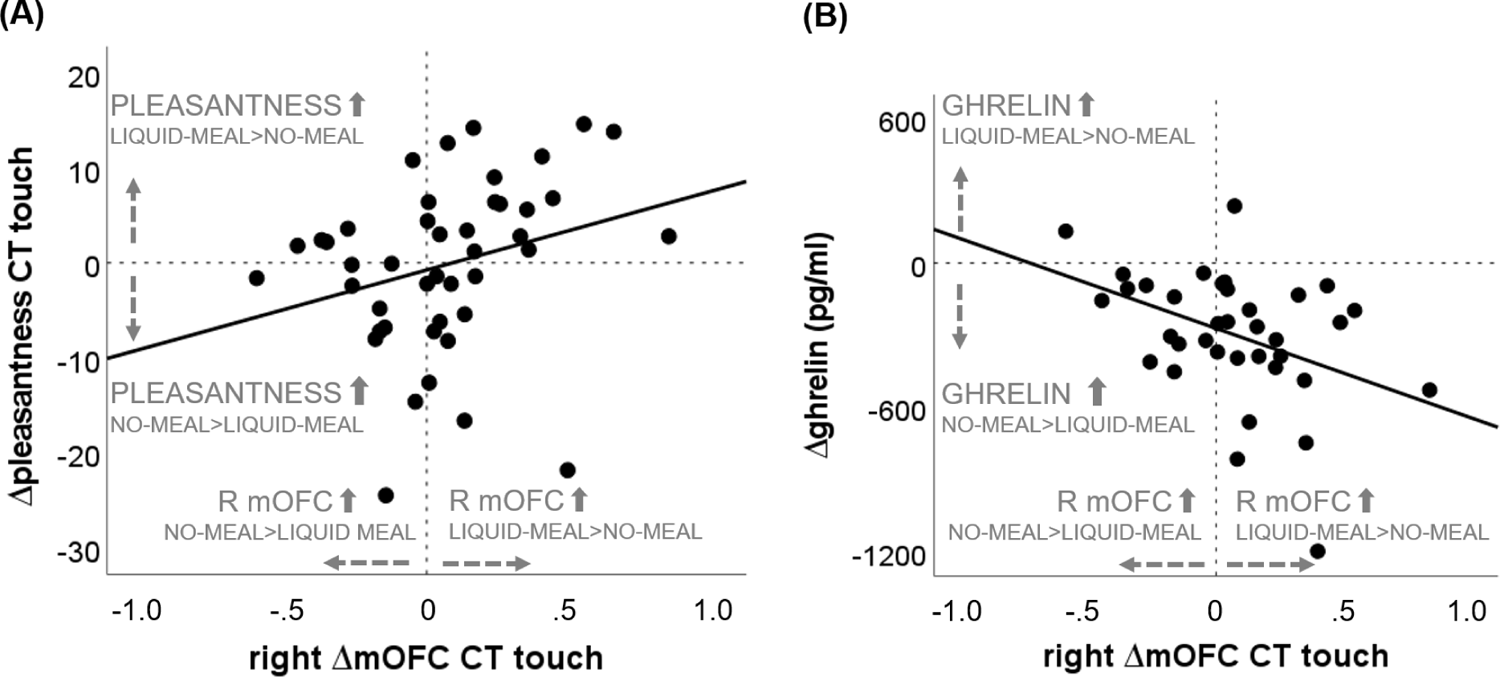
Nutritional state-dependent differences in brain activation and pleasantness ratings during CT-targeted touch and ghrelin variation. (A) “liquid-meal minus no-meal” session differences of right mOFC activation during CT-targeted touch on the x-axis (right ΔmOFC CT touch; positive values indicate higher right mOFC activation in the liquid-meal than the no-meal session; negative values indicate higher right mOFC activation in the no-meal than the liquid-meal session). On the y-axis, “liquid-meal minus no-meal” session differences in mean pleasantness ratings are depicted (Δpleasantness CT touch; positive values indicate higher reported pleasantness of CT-targeted touch in the liquid-meal than the no-meal session; negative values indicate higher reported pleasantness in the no-meal than the liquid-meal session). (B) “liquid-meal minus no-meal” session differences of right mOFC activation during CT-targeted touch on the x-axis (right ΔmOFC CT touch). On the y-axis, “liquid-meal minus no-meal” session differences in ghrelin concentrations at T1 are depicted (Δghrelin (pg/ml); positive values indicate higher ghrelin concentrations in the liquid-meal than the no-meal session; negative values indicate higher ghrelin concentrations in the no-meal than the liquid-meal session), this represents the satiety response induced by the meal resulting in ghrelin suppression in the liquid-meal session. The solid black lines depict the regression lines per correlation.

## 4. Discussion

This study investigated reward responses to touch after fasting participants received a liquid meal or no meal. Two main hypotheses were investigated: 1) Nutritional state and ghrelin modulate the rewarding effect of touch, measured as rated pleasantness and brain activation, and b) Differences in ROI activation during CT-targeted touch after versus without a meal would be associated with differences in ghrelin.

All touch was experienced as less pleasant without the meal while experienced pleasantness was not related to ghrelin concentrations. In contrast, the right medial orbitofrontal cortex (mOFC) was less activated during all touch when ghrelin concentrations were high. Thus, hypothesis 1 was partly supported by our findings. Hypothesis 2 was supported as a larger ghrelin decrease after the meal was associated with higher mOFC activation during CT-targeted touch, and this OFC activation was associated with higher experienced pleasantness.

### 4.1 Touch pleasantness was higher after the meal than without, and brain activation was higher after the meal than without

All touch velocities were rated as less pleasant when participants had not eaten. This may be because the participants may have been distracted by hunger-related interoceptive feelings caused by an empty stomach. Previous studies found that hunger states divert participants’ attention from a current task (Sänger, 2019; Stamataki et al., 2019). Another explanation might be that being hungry is often accompanied by an unpleasant subjective feeling in humans (MacCormack and Lindquist, 2019), which could have decreased the experienced pleasantness of touch. Indeed, participants reported slightly higher negative affect without a meal than following the liquid meal, but statistically, negative affect was not related to brain activation or pleasantness (see Supplementary Materials, section 3.6).

The finding that touch is less pleasant when hungry is novel. A related finding could be that fasters during Ramadan were more risk/loss averse in economic gambling tasks, suggesting that hunger can reduce the subjective value of monetary rewards (Rad and Ginges, 2017). Although the effect on touch pleasantness was small with only 2%, this might be different in more natural situations. This could have implications for a wide range of domains. For example, it may imply that touch sought for reasons of well-being, such as a relaxing massage, has a more positive effect after a meal than on an empty stomach. The reduced pleasantness of non-food rewards could also contribute to the difficulty of sticking to a weight loss diet.

Participants’ nutritional state also affected brain activation during touch and the associated subjective experience. Brain activation in the right inferior frontal gyrus and frontal operculum was higher following eating than fasting. As we had no hypothesis about these brain areas, this finding needs further investigation, and will not be discussed further.

### 4.2. mOFC activation during touch was related to ghrelin concentrations

Higher ghrelin concentrations were associated with reduced right mOFC activation in response to all touch velocities (exploratory analyses). In addition to reduced mOFC activation with higher ghrelin concentrations during all types of touch, there was also an activation change specific to CT-targeted touch. The larger the individual ghrelin suppression following the liquid meal (i.e., the satiety response induced by the meal), the higher the mOFC activation during CT-targeted touch (pre-registered correlation). Thus, both pre-registered and exploratory analyses suggest that ghrelin is related to mOFC activation in a way that high ghrelin concentrations go along with reduced mOFC activation during touch. mOFC has been associated with ascribing hedonic value to different types of stimuli (Li et al., 2016; Padoa-Schioppa and Assad, 2006; Rolls, 2020; Setogawa et al., 2019) including touch (Rolls et al., 2003). In accordance with this function, mOFC activation appears to represent reward valuation also in the present study: The difference in mOFC activation during CT-targeted touch after a meal versus no meal was positively associated with the respective difference in pleasantness. Since ghrelin was related to reduced mOFC activation and mOFC activation in turn was related to pleasantness, ghrelin seems to reduce rather than enhance touch reward.

This finding can be interpreted in line with the suggestion that ghrelin acts as a negative valence signal (Schéle et al., 2017). In this study, artificially increasing ghrelin concentrations led to a learned place avoidance in rodents. A potential mechanism underling this observation could be that ghrelin activates agouti-related peptide (AgRP) neurons in the hypothalamic arcuate nucleus (Betley et al., 2015), which are involved in energy balance processes. Rodents dislike the activation of these neurons, and they counteract it with motivated behaviours such as food search and feeding. These behaviours reduce the firing rate of AgRP neurons because ghrelin concentrations drop in response to feeding (Betley et al., 2015). Therefore, the activation of these neurons has been suggested to transmit a signal with negative valence that motivates (teaches) the rodents to search for food in order to “turn off” the signal. Applying this logic to human research, pairing high ghrelin concentrations with innocuous stimuli such as touch could result in a negative experience. Touch would become negative (or less positive) simply because it is paired with a negative valence signal, i.e. via classic conditioning.

Such a negative “tagging” may have contributed to the effect that all touch was rated as less pleasant without the meal than following a meal, although ghrelin concentrations were not directly related to pleasantness (discussed in next section). However, ghrelin is only one of the many factors that change in a fasted versus a fed state, and may alone not be sufficient to induce changes in conscious pleasantness experiences.

mOFC activation has previously been found to be related to ghrelin concentrations when participants looked at food compared to landscape pictures. During the food pictures, left mOFC activation was increased following exogenous ghrelin administration, but not following saline administration (Malik et al., 2008). Both this and the present findings can be reconciled by assuming that ghrelin may indicate metabolic stimulus salience. In other words, ghrelin may amplify the salience of stimuli that can restore homeostatic balance, and may dampen the salience of stimuli that do not. This function has been attributed to the hypothalamic hunger system, which is the primary target region for ghrelin (Cowley et al., 2003). Ghrelin might tune mesolimbic dopaminergic neurons to enhance or attenuate the salience of potential reward stimuli (Cassidy and Tong, 2017), for example by modulating their firing rate or synaptic activity. Providing a physiological basis for this assumption, studies in rodents have demonstrated that neural circuits of reward processing – in particular the mesolimbic dopamine system – and neural circuits of appetite in the hypothalamus are closely interconnected and interdependent (Abizaid, 2009; Cassidy and Tong, 2017). Applied to the current findings, ghrelin may down-regulate the salience of the non-food stimulus touch. This seems to be achieved by specifically down-regulating brain activation reflecting the value of touch, since experienced intensity and the associated brain activation were not related to ghrelin concentrations. Touch is not helpful in re-establishing homeostatic balance and may even be counterproductive if it distracts from food search. Thus, reducing the appeal of touch would help an individual to focus on foraging, which is essential for survival in a state of food deprivation. Indeed, both physiological hunger and a hunger-like state via the activation of AgRP neurons can suppress other motivated behaviours in favour of food seeking. Among these behaviours were water consumption, self-preservation and social interaction with conspecifics (Burnett et al., 2016) and prosocial behaviour in mice (Pozo et al., 2023). The authors interpreted these results in a way that energetic needs compete with other motivations and can therefore also control social and prosocial behaviour. Such an interpretation might also explain the results of the current study. For an organism it is important to constantly evaluate and prioritise multiple needs according to its state, context and opportunity. Recent evidence suggests that the appetite-suppressing hormone leptin may play such a role (Petzold et al., 2023), but ghrelin might be another mechanism through which different motivated behaviours are balanced.

Although speculative, ghrelin down-regulating the value of touch could have important implications for disorders that go along with high ghrelin concentrations. For example, individuals with anorexia who are characterized by chronically high ghrelin concentrations experience CT-targeted touch as less pleasant than healthy controls (Crucianelli et al., 2016). The current results suggest that ghrelin might be a mechanism that contributes to this effect. It would be interesting to find out if blocking ghrelin signaling, as is discussed to treat alcohol disorder (Farokhnia et al., 2019), increases the represented value of touch.

### 4.3 Experienced pleasantness was not related to ghrelin concentrations

As in many previous studies, subjective pleasantness ratings showed an inverted U-shaped curve with CT-targeted touch being rated as most pleasant (Croy et al., 2016; Gentsch et al., 2015; Hielscher and Mahar, 2017) on the group level, with expected inter-individual variations (Croy et al., 2021). Moreover, whole brain results as well as ROI analyses demonstrated that CT-targeted touch was accompanied by enhanced brain activation in somatosensory and social cognition areas, as previously reported (Morrison, 2016; Sailer et al., 2016).

In contrast to one part of the first hypothesis, ghrelin was not associated with explicit ratings of touch pleasantness, which may have several explanations. First, circulating ghrelin concentrations might not have been high enough to lead to measurable changes in participants’ subjective experience. Consistent with this, animal studies suggest that the effects of ghrelin could be dose-dependent. Systemic administration of very high doses of ghrelin induced avoidance behaviours in a place preference study (Lockie et al., 2015), while a lower dose induced preference behaviours in a similar experimental set-up (Jerlhag, 2008). Second, a number of additional processing steps take place in between neural activation in response to stroking and the subsequently given pleasantness ratings: participants need to bring touch experience to their conscious awareness, compare this experience to the presented rating scale, determine where on the presented scale this experience is best represented, prepare the response while remembering the decision and execute the motor response. Ghrelin’s effects might be limited to earlier stages of brain representation, whereas later stages of deciding and responding might be less sensitive to ghrelin’s influence. Third, as we observed changes in subjective pleasantness related to fasting, but not directly ghrelin concentrations, ghrelin alone may not be sufficient for changes in subjective experience.

Overall, it appears that functional imaging was able to reveal subtle associations of ghrelin that might otherwise have been missed if only explicit ratings would have been collected.

#### 4.4 Strengths and limitations of the study

One of the strengths of the present study was to induce and utilize intra-individual changes in ghrelin concentrations caused by fasting and eating. Most previous imaging studies compared intravenous ghrelin administration vs. placebo, and thereby induced short-lasting artificially high ghrelin concentrations that are often physiologically impossible (Kunath et al., 2016). Ghrelin concentrations after external administration were previously associated with enhanced activation in OFC in response to food pictures (Goldstone et al., 2014; Malik et al., 2008), which was interpreted as enhanced hedonic response to food cues. In contrast, with naturally varying ghrelin concentrations, the relationship between ghrelin concentrations and such hedonic reactivity to food cues was only modest (Wever et al., 2021). We postulate therefore that it would be more informative and ecologically more valid to investigate physiologically plausible intra-individual ghrelin concentrations than external administration in humans.

The study also had some limitations. One is that the applied within-subject fast-and-feed approach does not allow to infer causal relationships between ghrelin concentrations and touch reward. Another limitation is that power calculations were based on a medium-sized effect, and the final sample size was restricted by feasibility issues of functional imaging research. Moreover, the ratings of subjective bodily experience could have been taken shortly after each blood sample collection to better match subjective and objective markers of hunger.

## 5. Conclusion

Naturally circulating ghrelin concentrations after eight hours of fasting were associated with neural activation in response to touch in the right mOFC, a brain area involved in hedonic valuation. Higher ghrelin concentrations were associated with reduced mOFC activation, which in turn was related to lower pleasantness. Moreover, touch was rated as less pleasant and negative affect ratings were higher when participants were in a state of energy deficiency. In such a state, ghrelin might contribute to down-regulating the value of social stimuli to promote food seeking instead. Our results show that beyond its established role as an appetite-stimulating hormone, ghrelin is also involved in assigning value to social rewards such as touch.

## Data availability

All data, code, and materials will be made publicly available via the Open Science Framework after publiation: XXX Group-level T maps are available via https://neurovault.org/collections/12572/ The study was pre-registered on the Open Science Forum (https://osf.io/f9rkq). This manuscript was deposited at a preprint server bioRxiv (https://doi.org/10.1101/2022.05.10.491384) under a CC-BY-NC-ND 4.0 International license.

## Acknowledgements

We thank TINE BA for providing us with the products used as liquid meal for free. We thank Aiste Gvildyte Næss, Pietro Aleksander Rocco Berger Lapolla, Aleksandra Pusica, Thea Wiker Engelund, and Anbjørn Ree for their work with participant recruitment and data collection, and Federica Riva for helpful comments on a previous version of the manuscript.

## Author contributions

Original idea: Uta Sailer, Suzanne L. Dickson; Designed the study: Uta Sailer, Daniela M. Pfabigan, Bjørn S. Skålhegg; Provided input to methods: Erik Schéle; Performed the research: Erik R. Frogner, Daniela M. Pfabigan; Analysis of blood samples: Per M. Thorsby; Data curation: Daniela M. Pfabigan, Erik R. Frogner; Data analyses: Daniela M. Pfabigan; Data visualization: Daniela M. Pfabigan, Erik R. Frogner; Wrote the manuscript: Daniela M. Pfabigan, Uta Sailer; Provided comments on the manuscript: Suzanne L. Dickson, Bjørn S. Skålhegg, Erik R. Frogner; Acquired funding: Uta Sailer. All authors contributed and agreed to the final version of the manuscript.

