## Supplementary Materials for "Ghrelin is related to lower brain reward activation during touch"

### The «hunger» hormone ghrelin is associated with neural activation during touch, but not its affective experience

#### 1.) Detailed hypothesis as in the pre-registration

We pre-registered a bi-directional hypothesis stating that high ghrelin concentrations during fasting may either increase or decrease experienced and brain reward by touch. On the one hand, one could expect increased touch reward with high ghrelin concentrations, because ghrelin enhances reward-seeking behaviour in animals (1–3). In humans, alcohol craving is enhanced after external ghrelin administration (4), and positive mood following monetary rewards is greater after fasting (5). On the other hand, there is also reason to expect decreased touch reward with high ghrelin concentrations. Feelings of hunger may divert attention from the touch experience. Previous studies showed that hungry participants' attention was drawn away from the main task (6–8). A different reason for assuming decreased touch reward with high ghrelin concentrations is that ghrelin was suggested to carry a negative valence signal because it increased place aversion in rodents (9). In line with this, hunger is often associated with unpleasant feelings in humans (10). Such a negative valence signal is assumed to reinforce preferences for things that reinstate homeostasis, namely food (11). As touch does not help to restore metabolic homeostasis, touch reward may be reduced when participants are fasted and thus have high ghrelin concentrations.

#### 2.) Detailed information on Material and Methods

##### 2.1 Further information on participant demographics)

Handedness was assessed using the Edinburgh Handedness Inventory (12). Mean handedness score was 63.5 (SD: 55.6), with 53 participants being right-handed, five being left-handed, and eight being ambidextrous (n=2 missing). We refrained from only testing right-handed participants because we assumed no effects of lateralization on the results. Most participants were of self-declared European descent (n = 66), two of Asian descent.

Regarding highest education, 22.7% reported to have completed high school, 37.9% reported to have completed higher education of a 3-year duration, and 36.4% reported to have completed higher education longer than four years. The remaining 3% chose the answer format “other”.

As reported in the pre-registration, a priori power analysis for general ANOVA models (conducted in PANGEA) for the behavioural data recommended a sample size of 66 participants to reach a power of 80% for a middle-sized effect (d=0.45) of the interaction of touch velocity x nutritional state for the pleasantness ratings in a within-subjects design (3 [touch velocity: very slow, CT-targeted, fast] x 2 [nutritional state: liquid-meal, no-meal] within-levels, replicates=1) (13).

Output of the *a priori* power analysis conducted with PANGEA (13) with N=66 for the touch velocity x nutritional state interaction:

```
$total_sample_sizes
total_observations      Participants      replicates
                 396               66               1
$sources_of_variation
[1] P    N    T    P*N   P*T   N*T   P*N*T error
$power_results
noncentrality_parameter     degrees_of_freedom          power
                 2.830             130.000              0.802
```

*Note*: P = participants, M = nutritional state, M = touch velocity

We aimed to recruit 80 participants to account for potential dropouts, but the Covid-19 pandemic did not allow us to reach this goal. Deviating from the power analysis in the pre-registration, we applied single trial analyses for the behavioural data with linear mixed models instead of a means-based ANOVA design. Thus, the current behavioural analyses reached a power of 80% for a small effect of the touch velocity x nutritional state interaction (d=0.3, replicants=5 (i.e, 5 repetitions per touch velocity)) already with 36 participants.

Output of the single-trial power analysis conducted with PANGEA (13) with N=36 for the touch velocity x nutritional state interaction:

```
$total_sample_sizes
total_observations      Participants      replicates
                1080               36               5
$sources_of_variation
[1] P    N    T    P*N   P*T   N*T   P*N*T error
$power_results
noncentrality_parameter     degrees_of_freedom          power
                 2.850              70.000              0.803
```

Participants completed several questionnaires online and prior to arrival. In addition to the handedness inventory, relevant to this study are the Body Awareness Questionnaire (BAQ; (14), the Temporal Experience of Pleasure Scale (TEPS; (15), the Behavioral Inhibition System/Behavioral Activation System scale (BIS/BAS; (16), and the Social Touch Questionnaire (STQ; (17)).

Written informed consent was obtained from all participants prior to the experiment. The study was approved by the Regional Ethics Committee (REK South-East B, project 26699).

### 2.2 Further details regarding the experimental procedure

Upon arrival at the Institute of Basic Medical Sciences, University of Oslo, all participants signed the informed consent form and the functional magnetic resonance imaging (fMRI) security checklist. Each participant came in for two sessions on separate days – see Figure 1 in the main document for a graphical overview.

Shortly before the T1 blood and saliva samples, participants filled in questions about their subjective bodily states, such as subjective hunger feelings ("*How hungry are you right now?*"), feelings of stomach emptiness ("*How full is your stomach right now?*"), subjective thirst ("*How thirsty are you right now*?"), wanting to eat ("*How much do you want to eat food right now?*"), and capability to eat ("*How much food could you eat right now?*"). These questions were answered on a five-point Likert scale ranging from 1=*not at all* to 5=*very much*. Participants were further asked how much they would be willing to pay (in NOK) for their favourite food right now, and how much time had passed since their last meal. Further, they filled in the Positive and Negative Affect Schedule (PANAS; Watson et al., 1988) to assess their affective state. These questions and the PANAS were collected with nettskjema.no, a survey developed and hosted by the University of Oslo.

### 2.3 Saliva analysis

For saliva samples, Salivette® tubes (Sarstedt AG &Co, Germany) were used (60 seconds chewing of a cotton ball). The Salivettes were centrifuged for 2 minutes at room temperature and 1000g at the end of the testing session before storage at -80ºC.

Saliva samples were analysed at the Dresden LabService GmbH (Dresden, Germany). Salivary concentrations were measured using commercially available chemoluminescence immunoassays with high sensitivity (IBL International, Hamburg, Germany). The intra- and inter-assay coefficients for cortisol were below 9%.

### 2.4 Behavioural and hormone analyses

Linear mixed model descriptions for hormone analyses: hormones ~ 1 + nutritional_state + time_points + nutritional_state:time_points + (1 + nutritional_state | participant).

Touch ratings were also analysed with linear mixed models. We applied a two-stage approach to disentangle (i) effects of nutritional state per test session, and (ii) associations of ghrelin concentrations with subjective ratings during the task. In a first step, single-trial pleasantness and intensity ratings were modelled as a function of touch velocity (very slow/CT-targeted/fast), nutritional state (liquid-meal/no-meal), the mean-centred trial number, and the interaction of touch velocity x nutritional state as fixed effects. The random effects structure included a random intercept for participant and random slopes for touch velocity and nutritional state (model description: ratings ~ 1 + touch_velocity + nutritional_state + trial_number + touch_velocity: nutritional_state + (1 + touch_velocity + nutritional_state | participant). In a second step, single-trial pleasantness and intensity ratings were modelled as a function of touch velocity (very slow/CT-targeted/fast), measurement session (1,2), mean-centred trial number, and mean-centred ghrelin concentrations (sample time point T1 of each session), and the interaction of touch velocity x ghrelin as fixed effects. The random effects structure included a random intercept for participant and random slopes for touch velocity and measurement session (model description: ratings ~ 1 + touch_velocity + measurement_session + trial_number + ghrelin + touch_velocity:ghrelin + (1 + touch_velocity + measurement_session | participant).

### 2.5 fMRI analyses

Following pre-processing, single-subject analysis (first-level analysis) was performed based on the General Linear Model framework in SPM12 (19). Regressors of interest included one regressor for each touch velocity (very slow, CT-targeted, fast; duration: 15 sec), one regressor for each VAS scale (pleasantness, intensity; duration: individual response times), and one regressor-of-no-interest for the last five volumes of the scan session.

Deviating from the pre-registered analyses, we used two separate regressors for the two VAS scales (instead of one) and added a regressor-of-no-interest for the last five volumes because the scan session was longer than the presentation of the final fixation cross in a few participants. Trials with missed responses were not excluded from fMRI analysis because the neural response to touch preceded the response interval (those trials were excluded from behavioural analyses though). The regressors were convolved with the default canonical hemodynamic response function implemented in SPM12 (128 Hz high pass filter). fMRI pre-processing and first level analyses were performed on the high-performance computing resource Saga, owned by the University of Oslo, and operated by the Department for Research Computing at USIT, the University of Oslo IT-department, https://www.uio.no/english/services/it/research/hpc/saga/, running on MATLAB R2019a.

Group statistics ("second level analysis") was conducted in several steps. Using the 45 artefact-free participants, we first calculated a full factorial model (deviating from the pre-registration where we planned a random effects model) with touch velocity (very slow, CT-targeted, fast) and nutritional state (liquid-meal/no-meal). Results are presented at a cluster-level family-wise error (FWE) corrected threshold of $p<.05$ (starting threshold $p<.001$ uncorrected).

Second, we adopted a region-of-interest (ROI) approach. We pre-registered ROIs in somatosensory, social cognition, and reward processing networks. Based on recent literature three ROIs known to be involved in the processing of CT-targeted touch were pre-registered: left posterior insula, and left and right secondary somatosensory cortex (SII) (20–22). We expected enhanced activation for CT-targeted than for very slow and fast touch in these areas. To address multiple comparisons, results of the left posterior insula ROI and the two SII ROIs are presented at a Bonferroni-corrected level of p<.017 (.05/3 ROIs) per analysis. One ROI involved in social processing and cognition was pre-registered in the right superior temporal gyrus (STG) (23–25). We expected enhanced activation in STG for CT-targeted than for very slow and fast touch. For processing of the rewarding aspects of touch, two ROIs in medial orbitofrontal cortex (left and right mOFC) (26–28) were pre-registered (Bonferroni-corrected level of p<.025 (0.05/2 ROIs). We expected that CT-targeted touch would lead to enhanced mOFC activation compared to very slow and fast touch; and that ghrelin concentrations would be associated with mOFC activation.

Most of the ROIs were anatomically defined via the JuBrain Anatomy Toolbox v3.0 (29), (bilateral: SII: OP1, mOFC: Fo3; left-hemispheric: posterior insula: Ig2), because of the precise anatomical segregation of brain areas. A functional ROI was created for the right STG ROI using MarsBaR v0.43 SPM toolbox (spherical 10 mm ROIs centred at x/y/z = 63/-44/22, based on the study by (23).

Deviating from the pre-registration, we refrained from creating a hypothalamus ROI because the available scanning sequence was not optimized for this purpose. Instead, we explored activity in further brain reward areas (left and right ventral striatum (VS) ROIs; previously described in (30). In line with the mOFC hypothesis, we expected larger activation for CT-targeted than for very slow and fast touch, and an association with ghrelin. In addition, we explored brain activity in the interoceptive circuitry to assess whether nutritional state had an effect (left and right anterior insula (31); anatomical ROI Id7 from the JuBrain Anatomy Toolbox v3.0). We expected enhanced activation in the no-meal compared to the liquid-meal session reflecting enhanced interoceptive attention (32). Results of the exploratory ROI analyses are presented at a Bonferroni-corrected level of p < .013 (.05/4 ROIs) per analysis.

Parameter estimates were extracted from the ROIs using the REX toolbox (http://web.mit.edu/swg/software.htm).

For the ROI analyses with linear mixed models, 60 participants were available in the liquid-meal session and 51 participants in the no-meal session. Again, we applied a two-stage approach to disentangle effects of nutritional state and ghrelin concentrations on brain activation. Analogously to the behavioural analyses, mean ROI activation was first modelled as a function of touch velocity (very slow/CT-targeted/fast), nutritional state (liquid-meal/no-meal), and their interaction as fixed effects. The random effects structure included a random intercept for participant and a random slope for nutritional state; if model convergence allowed it a random slope for touch velocity was added (model description: ROI_activation ~1 + touch_velocity + nutritional_state + touch_velocity: nutritional_state + (1 + touch_velocity + nutritional_state│ participant). Second, mean ROI activation was modelled as a function of touch velocity (very slow/CT-targeted/fast), measurement session (1,2), and mean-centred ghrelin concentrations (sample time point T1 of each session), and the interaction of touch velocity x ghrelin as fixed effects. The random effects structure included a random intercept for participant and random slopes for measurement session and touch velocity (if model convergence allowed the latter) (model description: ROI_activation ~1 + touch_velocity + measurement_session + ghrelin + touch_velocity:ghrelin + (1 + touch_velocity + measurement_session│ participant).

As described in the pre-registration, in a last step, potential associations between differences in CT-targeted touch ROI brain activation following liquid-meal vs. no-meal and differences in nutritional state measures, in particular ghrelin, were investigated. We calculated Spearman correlations between difference values (liquid-meal minus no-meal) of ghrelin (labelled as Δghrelin) and ROI brain activation. To limit the number of conducted correlations, this was done only for the one ROI (right mOFC; labelled as ΔmOFC) showing an association of ghrelin with brain activation in the exploratory ROI analyses. In an exploratory manner, we also tested whether differences in pleasantness (Δpleasantness) and intensity (Δintensity) ratings for CT-targeted touch were associated with differences in right ΔmOFC (Bonferroni-corrected $p<.025$ (0.05/2 correlations).

### 2.6 Exploratory control analyses

Negative affective state ratings assessed with the PANAS differed significantly between the liquid-meal and the no-meal session. Therefore, we ran exploratory analyses to rule out effects of negative affect on ratings and brain activation. The following linear mixed models were conducted for pleasantness and intensity ratings (model description: ratings ~ 1 + touch_velocity + measurement_session + trial_number + ghrelin + PANAS_negative + touch_velocity:ghrelin + touch_velocity:PANAS_negative + ghrelin_leves:PANAS_negative + (1 + touch_velocity + measurement_session│participant) and ROI activation (model description: ROI_activation ~1 + touch_velocity + measurement_session + ghrelin + PANAS_negative + touch_velocity:ghrelin + touch_velocity:PANAS_negative + ghrelin_leves:PANAS_negative + (1 + touch_velocity + measurement_session│ participant).

Moreover, we calculated Spearman correlations to explore whether ghrelin concentrations (at T1) were associated with affective state ratings (PANAS).

### 3.) Detailed results description:

#### 3.1 Effects of fasting on bodily and affective states

In response to a reviewer comment, we explored whether ghrelin concentrations at T1 and subjective hunger ratings were associated with each other in the no-meal and the liquid-meal test session. Conducting Spearman correlations, we observed no significant relationship between subjectively experienced hunger and ghrelin concentrations at T1 in the liquid-meal ($r_s$=-.174, p=.196, N=57) and the no-meal condition ($r_s$=-.058, p=.681, N=52).

**Table S1. Effects of liquid-meal and no-meal on bodily and affective states**

| | **Liquid-meal** | | **No-meal** | | | |
|---|---|---|---|---|---|---|
| | *M* | *SD* | *M* | *SD* | *p* | *effect size* |
| **Blood glucose (mmol/L)** | 5,07 | 0,45 | 4,95 | 0,44 | .067 | -0.24[a] |
| **Bodily state ratings** | | | | | | |
| Hunger feeling | 2,34 | 0,97 | 3,15 | 1,01 | **< .001** | .784[b] |
| Full stomach | 2,98 | 1,11 | 1,50 | 0,62 | **< .001** | -.933[b] |
| Thirst | 1,63 | 0,93 | 2,63 | 0,95 | **< .001** | .948[b] |
| Wanting to eat | 2,48 | 1,11 | 3,52 | 1,08 | **< .001** | .807[b] |
| Capability to eat | 3,00 | 1,03 | 3,85 | 0,81 | **< .001** | .966[b] |
| **Willingness to pay for food** (in NOK) | 113,00 | 102,00 | 172,00 | 122,00 | **< .001** | .876[b] |
| **PANAS** | | | | | | |
| Positive Affect | 29,70 | 6,67 | 28,70 | 6,94 | .379 | -0.11[a] |
| Negative Affect | 11,80 | 2,68 | 12,70 | 3,30 | **.007** | .475[b] |

*Note:* a = Cohen's d; b = rank biseral correlation

### 3.2 Ghrelin and cortisol concentrations at the three measurement time points

Ghrelin variation (in pg/mL) is depicted in Figure S1.

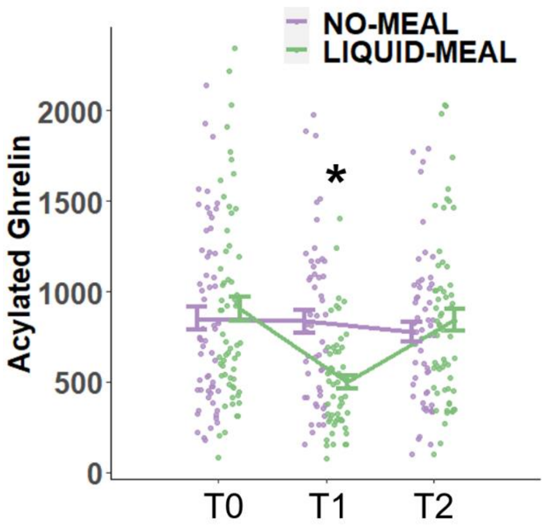


**Figure S1**: **Mean and individual acylated ghrelin concentrations** (pg/ml) for the three measurement time points in the no-meal (violet) and the liquid-meal (green) session. Error bars denote SEM. * denotes significant differences at T1 ($p<.001$).

Looking specifically at the liquid-meal session, ghrelin concentrations decreased from T0 (baseline at the beginning of the experiment) to T1 (shortly after the liquid meal) in all participants with blood samples from both measurement time points. Values ranged from 4 – 1651 pg/ml; M=384,21, SD=41.11; n=56. This was also indicated by a significant post-hoc test in the linear mixed model with ghrelin as outcome variable (difference =386.87, SE=29.1, t(225)=13.299, $p<0.001$).

For cortisol concentrations, we excluded two participants from further analyses due to implausible physiological values. Cortisol concentrations were only affected by sample time point. While T0/baseline and T1 concentrations did not differ from each other (b=0.14, SE=0.45, t(244.0)=0.32, p=.751), cortisol was significantly decreased at T2 compared to T1 (b=1.97, SE=0.45, t(244.0)=4.36, $p<.001$, semi-partial $R^2$=0.10). No other effects were significant (all p-values > .658). Means are depicted in Table S2.

**Table S2. Cortisol concentrations**

| | Liquid-meal session | | | No-meal session | | |
|---|---|---|---|---|---|---|
| | *M* | *SD* | *n* | *M* | *SD* | *n* |
| **Cortisol (nmol/l)** | | | | | | |
| *Baseline (T0)* | 5,02 | 4,51 | 64 | 4,95 | 3,31 | 60 |
| *T1* | 4,93 | 5,64 | 64 | 4,79 | 5,96 | 59 |
| *T2* | 3,08 | 2,63 | 64 | 2,67 | 2,43 | 60 |

### 3.3 Exploratory ROI based analysis of touch velocities, and their relationship to nutritional state and ghrelin concentrations

Higher ghrelin concentrations were associated with decreased brain activation in the right mOFC (b=-1.34$e^{-4}$, SE=5.45$e^{-5}$, t(90.2)=-2.46, p=.016, semi-partial $R^2$=0.06) – see Figure S2. Detailed results of the exploratory ROI analyses are presented in Table S3.

**Table S3. Exploratory ROI-based analysis**

| | | | *b* | *SE* | *df* | *t* | *p* |
|---|---|---|---|---|---|---|---|
| **LMM touch velocity x nutritional state** | | | | | | | |
| **Somatosensory ROIs (p<.0125)** | | | | | | | |
| | *posterior insula left* | n. s. | | | | | all p's > .030 |
| | ***SII left*** | **touch velocity (slow vs. CT-targeted)** | -0.18 | 0.04 | 214.0 | -5.02 | **< .001** |
| | | remaining conditions | | | | | all p's > .179 |
| | ***SII right*** | **touch velocity (slow vs. CT-targeted)** | -0.09 | 0.03 | 214.0 | 3.31 | **0.001** |
| | | **touch velocity (CT-targeted vs. fast)** | 0.16 | 0.03 | 214.0 | 5.56 | **< .001** |
| | | remaining conditions | | | | | all p's > .246 |
| **Social cognition ROI (p<.050)** | | | | | | | |
| | ***sup. temp. gyrus right*** | **touch velocity (slow vs. CT-targeted)** | -0.09 | 0.03 | 214.0 | -2.87 | **0.004** |
| | | remaining conditions | | | | | all p's > .077 |
| **Reward ROIs (p<.025)** | | | | | | | |
| | *mOFC left* | n. s. | | | | | all p's > .101 |
| | *mOFC right* | n. s. | | | | | all p's > .111 |
| **Exploratory ROIs (p<.0125)** | | | | | | | |
| | *VS left* | n. s. | | | | | all p's > .046 |
| | ***VS right*** | **touch velocity (slow vs. CT-targeted)** | -0.07 | 0.02 | 214.0 | -2.90 | **0.004** |
| | | remaining conditions | | | | | all p's > .055 |
| | *anterior insula left* | n. s. | | | | | all p's > .070 |
| | *anterior insula right* | n. s. | | | | | all p's > .078 |
| **LMM touch velocity x measurement session x ghrelin** | | | | | | | |
| **Somatosensory ROIs (p<.0125)** | | | | | | | |

| | | | | | | | |
|---|---|---|---|---|---|---|---|
| | *posterior insula left* | | | | | | all p's > .030 |
| | ***SII left*** | **touch velocity (slow vs. CT-targeted)** | -0.18 | 0.04 | 186.0 | -4.67 | **< .001** |
| | | remaining conditions | | | | | all p's > .029 |
| | ***SII right*** | **touch velocity (slow vs. CT-targeted)** | -0.09 | 0.03 | 186.0 | -2.74 | **0.007** |
| | | **touch velocity (CT-targeted vs. fast)** | 0.14 | 0.03 | 186.0 | 4.48 | **< .001** |
| | | remaining conditions | | | | | all p's > .077 |
| **Social cognition ROI (p<.050)** | | | | | | | |
| | ***sup. temp. gyrus right*** | **touch velocity (slow vs. CT-targeted)** | -0.10 | 0.03 | 186.0 | -3.03 | **0.003** |
| | | remaining conditions | | | | | all p's > .148 |
| **Reward ROIs (p<.025)** | | | | | | | |
| | *mOFC left* | n. s. | | | | | all p's > .108 |
| | ***mOFC right*** | **ghrelin** | -1.34e-4 | 5.45e-5 | 90.2 | -2.46 | **0.016** |
| | | remaining conditions | | | | | all p's > .712 |
| **Exploratory ROIs (p<.0125)** | | | | | | | |
| | *VS left* | n. s. | | | | | all p's > .054 |
| | ***VS right*** | **touch velocity (slow vs. CT-targeted)** | -0.07 | 0.03 | 186.0 | -2.86 | **0.005** |
| | | remaining conditions | | | | | all p's > .074 |
| | *anterior insula left* | | | | | | all p's > .014 |
| | *anterior insula right* | | | | | | all p's > .218 |

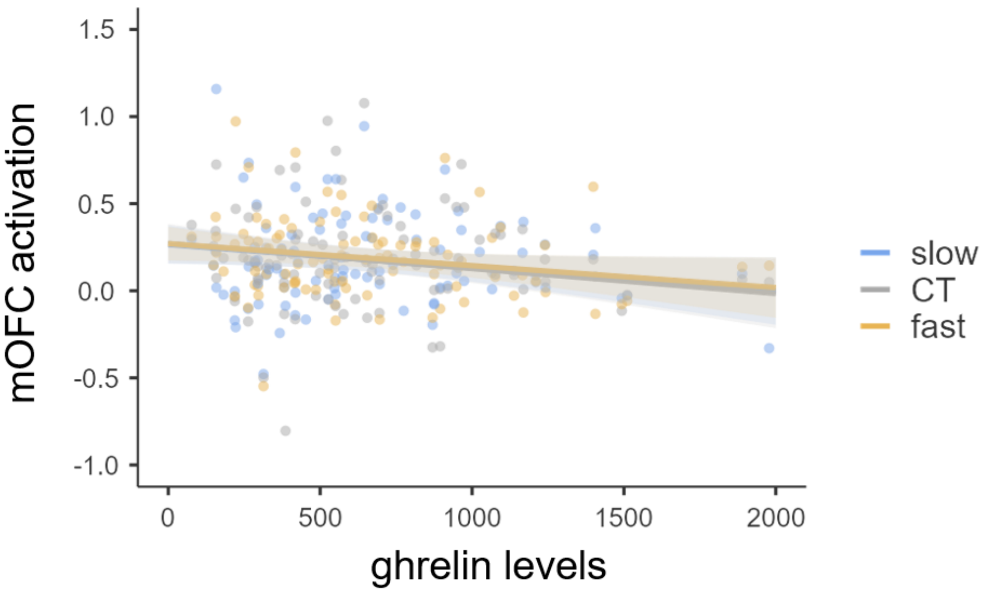


**Figure S2. Regression slope of ghrelin's association with right mOFC activation during touch.** The three touch velocities are depicted in different colours, the shaded area indicates a 95% confidence interval; ghrelin concentrations in pg/ml

### 3.4 Pre-registered relationship analysis between brain activation during CT-targeted touch and ghrelin concentrations

A significant negative association between Δghrelin and right ΔmOFC activation was observed ($r_s$=-0.412, p=.013, n=36). Further, a significant positive association between Δpleasantness and right ΔmOFC activation ($r_s$=0.377, p=.012, n=44) was observed. Importantly, Δghrelin and Δpleasantness were not associated with each other (p=.914). No significant correlations were observed for Δintensity with ΔmOFC activation or Δghrelin concentrations (both p-values > .516).

### 3.5 Exploratory analyses assessing the influence of PANAS negative affect on ratings and ROI activation, and on PANAS associations with ghrelin concentrations

No main effects of PANAS negative affect were found, see Table S4 for a detailed description.

No significant associations were observed between ghrelin concentrations and PANAS positive and negative affective state in the liquid meal (both p-values > .539) and the no meal session (both p-values > .657). Also session differences (liquid meal minus no meal) for ghrelin concentrations and PANAS positive and negative affect were not associated with each other (both p-values > .712).

**Table S4. Exploratory PANAS_negative analyses**

| | | | *b* | *SE* | *df* | *t* | *p* |
|---|---|---|---|---|---|---|---|
| **Ratings - LMM touch_velocity x measurement session x ghrelin x PANAS neg** | | | | | | | |
| **Pleasantness** | | | | | | | |
| | | PANAS negative | -0.078 | 0.447 | 105.3 | -0.18 | .862 |
| | | **touch (slow vs. CT)** | -11.718 | 1.347 | 60.9 | -8.70 | **<.001** |
| | | **touch (CT vs. fast)** | 4.557 | 1.590 | 58.0 | 2,87 | **.006** |
| | | **trial number** | -0.293 | 0.060 | 1425.6 | -4.92 | **<.001** |
| | | remaining conditions | | | | | all p's>.218 |
| **Intensity** | | | | | | | |
| | | PANAS negative | -0.260 | 0.513 | 106.1 | -0.51 | .613 |
| | | **touch (slow vs. CT)** | -10.399 | 1.544 | 58.9 | -6.74 | **<.001** |
| | | **touch (CT vs. fast)** | -14.673 | 2.243 | 60.5 | -6.54 | **<.001** |
| | | **trial number** | 0.336 | 0.062 | 1430.4 | 5,41 | **<.001** |
| | | **measurement session** | 3.542 | 1.372 | 54.8 | 2,58 | **.013** |
| | | remaining conditions | | | | | all p's>.107 |
| **LMM touch velocity x measurement session x ghrelin** | | | | | | | |
| **Somatosensory ROIs (p<.0125)** | | | | | | | |
| | *posterior insula left* | PANAS negative | -1.49e-5 | 0.014 | 81.5 | -0.001 | .999 |
| | | remaining conditions | | | | | all p's>.030 |
| | *SII left* | PANAS negative | 0.01 | 0.01 | 80.4 | 0.76 | .449 |
| | | **touch (slow vs. CT)** | -0.176 | 0.037 | 184.0 | -4.77 | **<.001** |
| | | **PANAS negative x touch (CT vs. fast)** | -0.046 | 0.015 | 184.0 | -3.01 | **.003** |
| | | remaining conditions | | | | | all p's>.036 |
| | *SII right* | PANAS negative | -0.009 | 0.011 | 85.3 | -0.83 | .411 |
| | | **touch (slow vs. CT)** | -0.087 | 0.031 | 184.0 | -2.77 | **.006** |
| | | **touch (CT vs. fast)** | 0.142 | 0.031 | 184.0 | 19815,00 | **<.001** |
| | | remaining conditions | | | | | all p's>.014 |

| | | | | | | | |
|---|---|---|---|---|---|---|---|
| **Social cognition ROI (p<.050)** | | | | | | | |
| | *superior temporal gyrus right* | PANAS negative | -4.26e-4 | 0.013 | 80.4 | -0.033 | .973 |
| | | **touch (slow vs. CT)** | -0.101 | 0.033 | 184.0 | -3.05 | **.003** |
| | | **PANAS negative x touch (slow vs. CT)** | 0.030 | 0.014 | 184.0 | 2,21 | **.028** |
| | | remaining conditions | | | | | all p's>.087 |
| **Reward ROIs (p<.025)** | | | | | | | |
| | *medial OFC left* | PANAS negative | -0.005 | 0.009 | 83.0 | -0.53 | .598 |
| | | **PANAS negative x touch (CT vs. fast)** | -0.026 | 0.010 | 184.0 | -2.71 | **.007** |
| | | remaining conditions | | | | | all p's>.105 |
| | *medial OFC right* | PANAS negative | -0.016 | 0.009 | 82.6 | -1.59 | .115 |
| | | **ghrelin** | -1.25e-4 | 5.48e-5 | 90.2 | -2.29 | **.024** |
| | | remaining conditions | | | | | all p's>.025 |
| **Exploratory ROIs (p<.0125)** | | | | | | | |
| | *VS left* | PANAS negative | -0.008 | 0.009 | 78.9 | -0.85 | .397 |
| | | **PANAS negative x touch (slow vs. CT)** | 0.027 | 0.011 | 184.0 | 2,61 | **.010** |
| | | **PANAS negative x touch (CT vs. fast)** | -0.29 | 0.011 | 184.0 | -2.74 | **.007** |
| | | remaining conditions | | | | | all p's>.050 |
| | *VS right* | PANAS negative | -0.011 | 0.009 | 80.8 | -1.17 | .247 |
| | | **touch (slow vs. CT)** | -0.071 | 0.025 | 184.0 | -2.89 | **.004** |
| | | remaining conditions | | | | | all p's>.024 |
| | *anterior insula left* | PANAS negative | 0.011 | 0.011 | 84.1 | 1,01 | .315 |
| | | remaining conditions | | | | | all p's>.017 |
| | *anterior insula right* | PANAS negative | -0.005 | 0.011 | 81.9 | -0.42 | .675 |
| | | remaining conditions | | | | | all p's>.212 |

### 3.6 Exploratory analyses with sex and age as covariates

In response to a reviewer's comments, we have computed exploratory analyses assessing the effects of participants' biological sex and age on the current results. Covariates such as sex and age were not considered in the preregistered hypotheses because the planned sample size would not have been large enough to investigate their effect with sufficient statistical power. In addition, the current sample with about 1/3 women is not well-balanced regarding sex. The following findings should therefore be interpreted with caution.

Previous studies reported sex differences in plasma ghrelin concentrations (see for example (33). Moreover, one study also reported an age-dependent decline in circulating acylated ghrelin concentrations in a fasted state (34). Moreover, there is further evidence that the subjective experience of social touch is related to one's age (35) and sex/gender (36). Based on the available literature, we tested whether ghrelin concentrations and subjective ratings would be influenced by sex and age.

#### Impact of sex and age on ghrelin concentrations

We added sex (women/men) and age as covariates to the linear mixed model, by modeling both their main effects on ghrelin concentrations. While age had no influence on ghrelin concentrations ($p=.401$), the covariate sex was significant ($p<.001$). We therefore added interaction terms of sex x nutritional state, sex x sample time point, and their three-way interaction to the model, which improved the model fit. A significant main effect of sex was found ($b=450.6$, $SE=109.7$, $t(61.5)=4.05$, $p<.001$, semi-partial $R^2=0.22$). Women had higher ghrelin concentrations than men, which is in line with the literature (e.g., (33)). There was also a significant three-way interaction for the comparison of the baseline timepoint and T1 ($b=310.2$, $SE=90.8$, $t(222.5)=3.42$, $p<.001$, semi-partial $R^2=0.05$). Descriptively resolving the three-way interaction, it looks as if women displayed a steeper ghrelin suppression caused by the meal than men (see right panel in the figure below, change from T0 to T1). In contrast, without the meal, only the main effect of sex (with higher concentrations in women than men) was observable.

#### Impact of sex and age on pleasantness and intensity ratings

To control for age and sex effects on pleasantness and intensity ratings, we added both variables as covariates to the respective linear mixed models as fixed effects and modelled them as main effects. No significant effects of participants' sex or age were observed for pleasantness ratings (nutritional state x touch velocity x trial number model: age: $p=.268$, sex: $p=.659$; ghrelin x measurement timepoint x touch velocity x trial number model: age: $p=.438$, sex= $p=.609$) or for intensity ratings (nutritional state x touch velocity x trial number model: age: p=.777, sex: p=.402; ghrelin x measurement timepoint x touch velocity x trial number model: age: p=.478, sex= p=.529).
